# When parasites bite hardest: mistletoe effects on oak radial growth peak near climatic optima

**DOI:** 10.1101/2025.08.24.671974

**Authors:** Jiří Doležal, Vojtěch Lanta, Kirill Korznikov, Lenka Plavcová, Jan Tumajer

## Abstract

Hemiparasitic mistletoes can alter host water and carbon balance, but their impact on tree growth is expected to vary with phenology, microclimate and stand context. We asked whether the yellow mistletoe *Loranthus europaeus* shifts the timing or reduces the magnitude of radial growth in *Quercus robur*, and whether any penalty is strongest near climatic optima for host growth. We instrumented 34 mature oaks across age (young, old), canopy position (solitary, closed-canopy), and infection (infected, non-infected) with point dendrometers at 15-minute intervals for four growing seasons (2020–2023). Air and soil temperatures and soil moisture were logged concurrently. We derived growth phenology, difference curves (non-infected minus infected), monthly climate–growth correlations, and response surfaces in temperature– moisture space. Growth phenology was consistent among years: onset around day-of-year 120–140, peak 150–220, cessation 250–270. Mistletoe did not shift onset or cessation but reduced growth amplitude, especially in high-growth years and solitary, well-lit trees. Suppression was greatest near climatic optima (≈10–18 °C with adequate soil moisture) and diminished when conditions were suboptimal (hot and dry or cold and wet), so infected and non-infected converged. Short-term climate–growth relationships were similar across infection status: temperature effects were negative during the main season, whereas soil moisture effects were positive. Young, solitary, non-infected trees responded more to mid-summer moisture than infected trees, consistent with infection shifting hosts from resource-tracking to stress-limited growth under exposure. Joint temperature–moisture response surfaces for infected versus non-infected trees were highly similar, indicating that mistletoe reduces growth magnitude, not niche. Our results identify the environmental window in which host–parasite competition bites hardest and provide a baseline for forecasting parasite impacts under shifting temperature and moisture regimes. Because the largest penalties arise near growth optima, the frequency of cool, moist periods may modulate impacts at the stand scale, particularly for solitary oaks at woodland–grassland ecotones. Integrating fine-scale growth with microclimate clarifies when hemiparasites depress performance and why effects vary across years, sites and canopy contexts. Results underscore the value of continuous dendrometer records for quantifying parasite impacts.

## Introduction

Parasitic plants are a distinctive and ecologically important component of terrestrial ecosystems, influencing the physiology, growth, and survival of their host plants while also affecting community dynamics and ecosystem functioning (Cameron et al., 2008; Mathiasen et al., 2008). Among these, mistletoes are hemiparasites that photosynthesize but rely on their hosts for water and mineral nutrients (Urban et al., 2012), often exerting significant impacts on host carbon balance, hydraulic status, growth and reproductive output (Watson, 2001; Press and Phoenix, 2005). By altering host performance, mistletoe infection can also influence competitive interactions and forest structure, making it an important driver of vegetation dynamics in many woodlands (Matula et al., 2015; Oladi et al., 2025; Gosling et al., 2024).

The effects of mistletoe on host trees, however, are highly context dependent. Variation in host susceptibility, canopy position, tree age and size, and local environmental conditions can mediate the magnitude and even the direction of infection effects (Zakaria et al., 2014; Santiago-Rosario et al., 2023). Climatic conditions may modulate host–parasite interactions, as both partners’ physiological activity responds non-linearly to temperature and water availability (Glatzel, 1983). Mistletoe impacts may vary not only among sites but also within a growing season, with potential seasonal shifts linked to host phenology and climatic optima for growth (Cocoletzi et al., 2020). Seasonal shifts in nutrient profiles in host and mistletoe xylem sap illustrate fluctuating interaction dynamics (Escher et al., 2004). Moreover, drought exacerbates growth decline in mistletoe-infested hosts (Tamudo et al., 2021), and mistletoe’s failure to close stomata under dry periods further stresses the host (Walas et al., 2022). Ongoing climate variability magnifies these resource imbalances, altering carbon, water, and nutrient dynamics in infected trees (González de Andrés et al., 2024).

In oaks, early-season radial growth often begins before leaf flush, when transpiration is minimal (Pérez-de-Lis et al., 2016; Lavrič et al., 2017; Szatniewska et al., 2022; Puchałka et al., 2024). This phenological timing raises the possibility that differences between infected and non-infected trees may be smaller during the initial growth phase and more pronounced later in the season, when leaves are fully expanded, and transpiration demand is high. Furthermore, differences in growth appear to be greatest near the host’s climatic optimum, when absolute growth rates are highest, and to diminish under suboptimal conditions such as extreme temperatures or low soil moisture, consistent with climate-modulated parasite impacts on hosts (Bell et al., 2019; Tamudo et al., 2021). This pattern accords with host–parasite theory for hemiparasites: under favourable conditions, both host and parasite exhibit high physiological activity, intensifying resource competition, whereas under stressful conditions the parasite may rely more heavily on its own photosynthesis (Press & Phoenix, 2005; Těšitel, 2016), with seasonal shifts in xylem solutes and stomatal behaviour further modulating outcomes (Escher et al., 2004; Walas et al., 2022). However, exceptions can occur, such as in shaded young trees, where infection effects may be reversed, suggesting complex interactions between microhabitat, host vigour, and parasite strategy (Queijeiro-Bolaños et al., 2016; Lázaro-González et al., 2019).

Recent studies reinforce this view of context-dependent outcomes. Queijeiro-Bolaños et al. (2016) showed that the balance between competition and facilitation in dwarf mistletoe– pine systems shifts with host size, stand density, and environmental stress, with infection effects sometimes neutral or even positive under resource-limited conditions. Lázaro-González et al. (2019) further demonstrated that mistletoes can generate non-trophic, trait-mediated effects that propagate through ecological networks, altering both host physiology and interactions with other organisms. These findings suggest that mistletoe impacts on radial growth cannot be understood solely in terms of resource extraction, as they must also be considered in the context of environmental variation, seasonal host physiology, and indirect ecological effects.

*Quercus robur* and related oak species are foundational components of Central European forests, providing habitat and resources for a wide range of organisms (Lanta et al., 2025). They are also known hosts for the European mistletoe (*Loranthus europaeus*), whose infection prevalence can be high in certain stands (Matula et al., 2015; Dolezal et al., 2010). Despite the recognized importance of this parasitic association, relatively few studies have examined its effects on host radial growth at fine temporal resolution, and even fewer have considered how these effects might vary seasonally or in relation to climatic drivers (Dolezal et al., 2016). Such information is critical for understanding how parasitic plants interact with their hosts under current and future climates, particularly in the context of warming and shifting precipitation patterns (Tumajer et al., 2022).

High-resolution dendrometer measurements offer a powerful means to track tree growth dynamics in unprecedented detail, capturing subtle variations associated with infection, phenological stage, and microclimatic conditions (Zweifel et al., 2016; Deslauriers et al., 2007). Although widely applied to study seasonal rhythms and climate–growth relationships, they have not previously been used to examine the impacts of mistletoe infection on host tree growth. By combining multi-year continuous growth monitoring with microclimatic data, it becomes possible to disentangle the overall effect of mistletoe infection from its seasonal modulation and climatic sensitivity. High-precision electronic dendrometers, already well established in studies of boreal, temperate, and tropical trees (Worbes, 1995; Etzold et al., 2022; Plavcova et al., 2025), record radial stem size with sub-hourly resolution, enabling the assessment of growth rates from hourly to annual scales and the characterization of tree water storage (De Swaef et al., 2015; Salomón et al., 2022). Recent applications have shown that growth in temperate trees occurs primarily at night when turgor conditions favour cell expansion (Zweifel et al., 2021), while at seasonal scales, growth is restricted to favourable days within a broader climatic window (Etzold et al., 2022; Tumajer et al., 2022). These insights illustrate the power of dendrometer data for identifying how environmental drivers constrain tree growth, and here, we extend this approach to the novel context of mistletoe–host interactions.

In this study, we monitored radial growth of oak trees with and without mistletoe infection over four consecutive growing seasons (2020–2023) using high-resolution dendrometers. We examined how infection effects on radial tree increment varied with tree age, canopy position, and year-to-year climatic conditions, and whether these effects were modulated by seasonal phenology or by specific combinations of temperature and soil moisture. We also investigated whether infection effects were most pronounced near climatic optima for growth, as predicted by host–parasite theory for hemiparasitic plants. By integrating growth phenology, climate–growth relationships, and infection effects, our work provides new insights into the context-dependent nature of mistletoe–oak interactions in temperate woodlands.

## Material and Methods

### Study area

The research was conducted in the National Nature Reserve (NNR) Čertoryje, located in the White Carpathians Protected Landscape Area along the Czech–Slovak border (Fig. 1). The region forms part of a hilly chain (maximum altitude 970 m a.s.l.) composed of base-rich flysch sediments. The landscape is a mosaic of villages in narrow valleys, steep slopes covered by deciduous woodland, and grasslands on shallow slopes and plateaus. Traditional land use, preserved until the early 1980s due to the late arrival of communist-era land consolidation, has maintained a fine-scale habitat mosaic of meadows, pastures, small fields, orchards, and woodlots (Dolezal et al., 2010). The climate is temperate with a mean annual temperature of 8.8 °C (1900–2024 average) and mean annual precipitation of 699 mm (reference). Grasslands cover roughly 20% of the protected area, dominated by species-rich “Carpathian meadows” with exceptionally high plant diversity (Albert et al., 2019). The studied oak stands consist predominantly of *Quercus robur* interspersed with other broadleaved species such as rowan trees (*Sorbus aucuparia*, *Sorbus torminalis*), maples (e.g., *Acer platanoides*) and hawthorns (*Crataegus* sp.), and exhibit varying degrees of European mistletoe (*Loranthus europaeus*) infestation.

**Figure 1.**
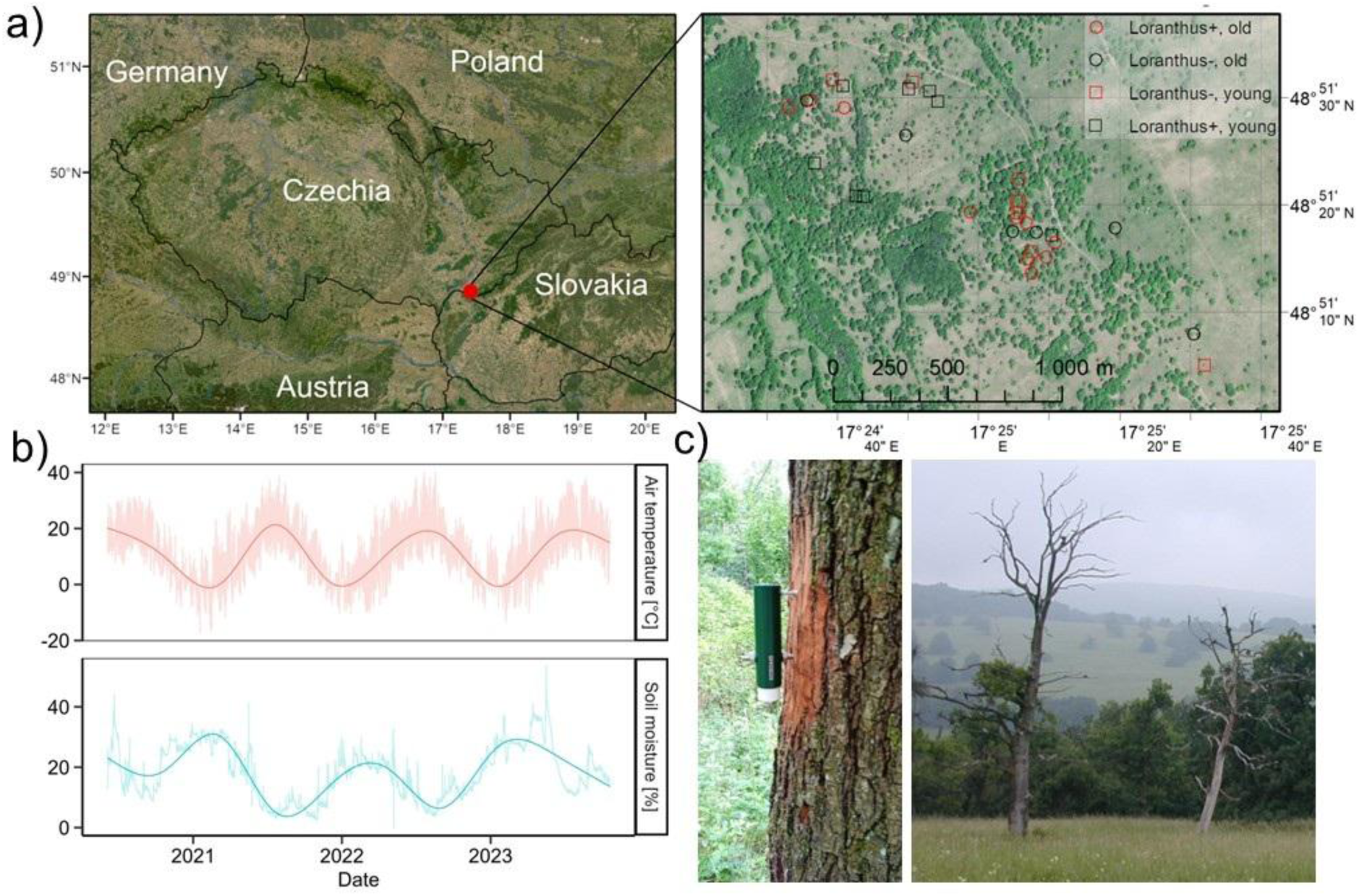
Study area (White Carpathians, Czech Republic). Savannah-like species-rich meadow with scattered oak groves and solitary trees at the woodland–grassland ecotone on base-rich flysch (a). High-resolution satellite map of the site showing mapped oaks by mistletoe (*Loranthus europaeus*) status and age class: red = parasitized (*Loranthus+*), black = unparasitized (*Loranthus–*); circles = old trees, squares = young trees. b) Site microclimate (2021–2023): air temperature (°C) and volumetric soil moisture (%), raw readings with a smoothed seasonal trend, both showing strong seasonality, warm, dry summers and cool, wetter winters. c) Left: Point dendrometer installed on an oak stem for continuous growth monitoring. Right: open woodland with severely declined/standing dead oaks illustrating canopy dieback in the landscape.

### Experimental design

We sampled 34 mature Quercus robur trees (Fig. 1a) covering eight combinations of age class (young, old), canopy position (solitary, closed-canopy), and mistletoe status (infected, non-infected). We aimed for four trees per combination (32 total), with two additional trees in the old × solitary × infected group (n = 6), which were instrumented with point dendrometers (TOMST, Praha, Czechia). Age classes were defined based on trunk diameter at breast height (DBH, smaller or larger than 30 cm) and number of growth rings at DBH counted on increment cores extracted using Pressler’s borer (younger or older than 60 years), and growth form: “young” trees were relatively vigorous; “old” trees often showed signs of age-related decline such as the presence of dead crown branches. Solitary trees were free-standing individuals exposed to full light, whereas canopy trees grew within closed stands and experienced greater shading. Infected trees bore visible mistletoe clumps, with infection intensity ranging from moderate to high. The number of mistletoe plants on each of the target trees with dendrometers was counted; infested trees had more than five plants (up to a maximum of 15 plants on a single tree (Doležal et al., 2016).

### Dendrometer installation and measurements

On each tree, a high-resolution point dendrometer (TOMST, Prague) was installed at ∼1.3 m height on the north-facing side of the stem to minimise direct solar heating. The installation took place in April 2020. The dendrometers recorded stem radius changes at 15-minute intervals throughout four consecutive growing seasons (2020–2023). Instruments were checked regularly to ensure stability and accuracy, and the final recorded data were downloaded during April 2023.

### Microclimatic measurements

Air temperature, soil temperature, and soil volumetric water content were monitored continuously using a network of TMS T4 sensors (Wild et al., 2019) positioned within the study area. Air temperature was measured at 15 cm above ground under the target trees, while soil temperature and moisture sensors were placed at 10 cm depth in representative microsites. All variables were logged at 15-minute intervals and synchronised with dendrometer data.

### Data processing

Raw electric signal of stem size variation (recorded by dendrometers) and soil moisture (TMS T4) was converted into radius increment (µm) and volumetric soil water content (%) using the ‘myClim’ package (Man et al., 2023) in R (R Core Team, 2024). Converted dendrometer data were visually checked for spurious values caused by mechanical disturbance or sensor drift. We used a zero-growth approach to calculate radial growth rate (µm h⁻¹), i.e., a positive difference between the instant stem radius and the previous stem radius maximum (Zweifel et al., 2017). During the periods of stem shrinking with the instant stem radius below the previous maximum, the radial growth rate was zero. We removed rare, unrealistic values of radial growth rate exceeding 150 µm per single timestamp (15 minutes) from further analysis. Mean daily growth curves were derived for each tree from radial growth rates at 15-minute resolution. Seasonal growth curves highlighting overall growth patterns beyond daily variation were represented by smoothing splines fitted to daily curves. Both daily and seasonal curves were aggregated into category means for further analysis.

### Statistical analyses

We analysed radial growth dynamics at three complementary scales. (1) Seasonal growth phenology - we identified the onset, peak, and cessation of growth for each category and year. Seasonal growth curves were visually compared between infected and non-infected trees to detect phenological shifts and amplitude differences. (2) Seasonal infection effects – To quantify infection effects over the course of the season, we calculated the difference in mean daily growth rates between non-infected and infected trees within each category (non-infected minus infected). Positive values indicated faster growth in non-infected trees. (3) Climate– growth relationships – Pearson correlation coefficients between 15-min radial growth rates and microclimatic variables (air temperature, soil temperature, soil moisture) were computed for each tree and calendar months, then averaged within categories. We also calculated mean radial growth rates across discrete climatic intervals (4 °C air temperature bins, 5% soil moisture bins) to identify climatic optima and assess whether infection effects (i.e., differences of mean growth rates between categories of infection) were strongest under optimal or suboptimal conditions. (4) Joint climatic effects – To visualise combined temperature–moisture effects, we plotted a matrix of mean growth rates for specific combinations of air temperature and soil moisture, with point size representing the frequency of occurrence and point colour indicating mean growth rate. These analyses were performed separately for infected and non-infected trees to identify differences in optimal climatic niches. We used the Mantel test to statistically evaluate differences between matrices for infected and non-infected trees. All data processing and statistical analyses were conducted in R v.4.2.2 (R Core Team, 2024) using packages dplyr and lubridate for data manipulation, myClim for pre-processing of dendrometer and climatic data, ggplot2 for plotting, base stats for correlation analysis, and vegan for the Mantel test. Figures were prepared with consistent axis ranges and colour scales to facilitate direct comparison among years and categories.

## Results

### Seasonal Growth Dynamics

Seasonal growth dynamics of oak trees, assessed over four consecutive growing seasons (2020–2023), revealed a consistent phenological pattern across all combinations of age class, canopy position, and mistletoe infection status (Fig. 2). Radial growth typically began between day of year (DOY) 120–140, peaked in late spring to early summer (DOY 150–220), and declined toward the end of the growing season (DOY 250–270). This pattern was stable among years and categories, with variation occurring mainly in the timing and magnitude of peak growth. Mistletoe infection did not alter the onset or cessation of growth in any year, but its effects were sometimes apparent in the amplitude of growth, particularly in favourable years.

**Figure 2.**
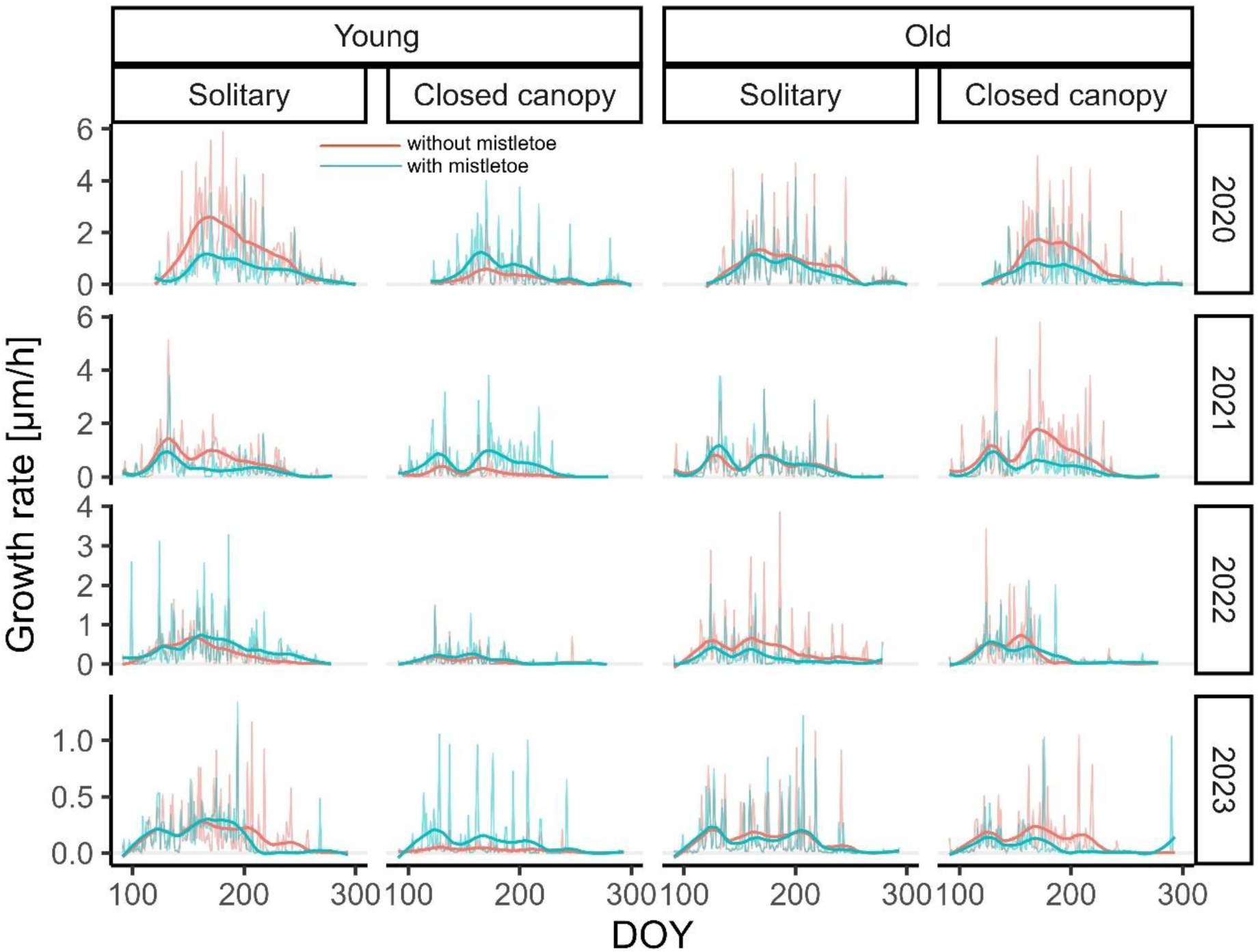
Daily and seasonal dynamics of radial growth rate (µm h^-1^) in oak trees from 2020 to 2023 between days of year (DOY) 90 and 300, shown by age class, canopy position, and mistletoe infection status. Red and blue lines indicate non-infected and infected trees, respectively; thin lines show mean daily growth rates, and thick lines show smoothed seasonal curves. Note different scales of y-axes between years.

### Interannual and Category-Specific Variation

In 2020, the year with the highest growth rates, young solitary trees reached seasonal peaks of growth rate 2–2.5 µm h⁻¹, with slightly reduced amplitudes in infected individuals. Old solitary and old canopy trees attained intermediate peaks (1.5–2 µm h⁻¹), while young canopy trees peaked at only ∼1–1.5 µm h⁻¹. In 2021, peak growth was lower (1.5–2 µm h⁻¹ in young solitary and old canopy trees; <1 µm h⁻¹ in other categories) and the active growth period appeared shorter, as indicated by narrower curves. Notably, growth was bimodal in 2021 with growth quiescence observed in all categories around DOY 150. In 2022, maximum growth declined further, with young solitary trees peaking at ∼0.8 µm h⁻¹ and all other groups at ∼0.4–0.6 µm h⁻¹. By 2023, growth rates were uniformly low (<0.5 µm h⁻¹) across all categories.

The influence of mistletoe infection on growth amplitude varied among years and tree categories. In high-growth years (2020–2021), infected trees often exhibited slightly lower seasonal and daily growth peaks, particularly among young solitary trees. However, in some cases, such as young canopy trees in all years and young solitary trees in 2022, infected individuals showed marginally higher average growth rates than uninfected ones. In low-growth years (2022–2023), differences between infection statuses were negligible, indicating that climatic constraints likely outweighed any parasitic effects.

Across all years, solitary trees consistently grew faster and exhibited greater variability than canopy trees, particularly in young trees. Overall, mistletoe infection did not influence the phenology of radial growth but affected growth magnitude in a context-dependent manner, with the direction and strength of the effect varying by year, age class, and canopy position.

### Seasonal Patterns of Infection Effects

Difference curves (non-infected minus infected; Fig. 3) revealed that positive values, indicating faster growth in non-infected trees, were most pronounced in 2020, especially in young solitary trees during the early-to-mid growing season (DOY 150–180; up to ∼1.5 µm h⁻¹). Old canopy trees in 2020 also displayed moderate positive peaks (∼0.75–1.0 µm h⁻¹). In 2021, differences were smaller and more transient, with occasional slight advantages for infected individuals in certain categories. In 2022, all categories showed differences mostly within ±0.3 µm h⁻¹, while in 2023 differences were minimal (<0.2 µm h⁻¹) and non-systematic. No consistent seasonal shift in infection effects was detected, suggesting that any influence of mistletoe on radial growth is not strongly tied to specific phenological phases of oak growth. The observed impacts are primarily context-dependent, emerging most clearly in high-growth years and varying according to tree age and canopy position.

**Figure 3.**
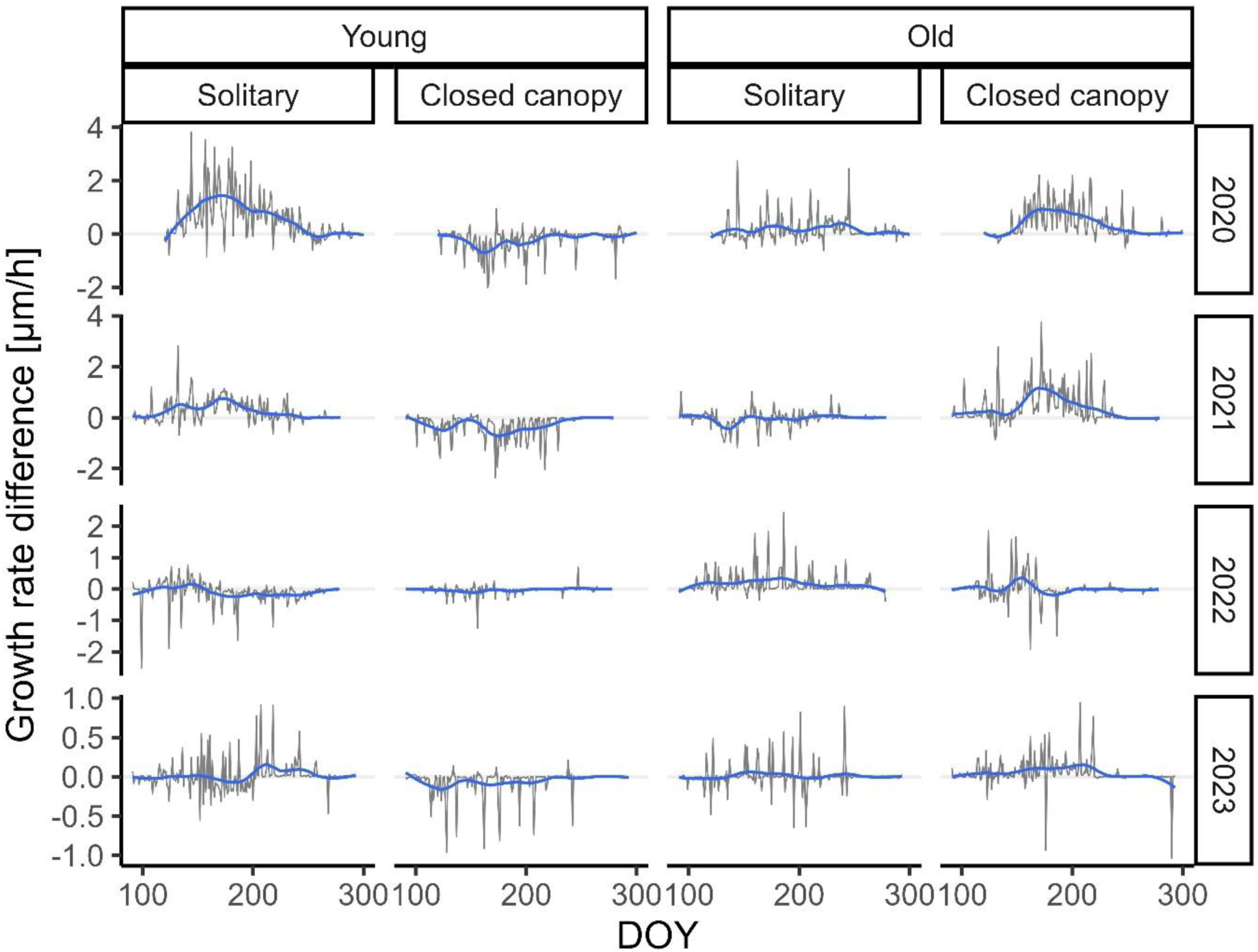
Daily and seasonal differences in radial growth rates between non-infected and mistletoe-infected oaks (non-infected minus infected) across age classes and canopy positions between days of year (DOY) 90 and 300 in calendar years 2020–2023. Positive values indicate faster growth in non-infected trees. Grey lines are based on mean daily growth rates; blue lines represent their seasonal smoothing splines. Note the different scales of the y-axis between years.

### Climate–growth relationships

Monthly correlations between radial growth rate (15-min resolution) and climatic variables (air temperature, soil temperature, and soil moisture) revealed significant associations across all age classes, canopy positions, and infection statuses (Fig. 4). For both air and soil temperature, correlations were mostly negative during the main growing season (May–August), indicating reduced growth under warmer conditions. Patterns were broadly consistent across years, with no systematic differences between infected and non-infected trees. Soil moisture generally showed significant positive correlations with growth, especially in early summer (June–July) for young solitary and old solitary trees. A notable pattern emerged in young solitary oaks: the mid-summer moisture dependence was stronger in non-infected trees than in those infected by mistletoe. Overall, the data suggest that short-term growth responses to microclimatic variation are modest, with soil moisture exerting more positive influence than temperature.

**Figure 4.**
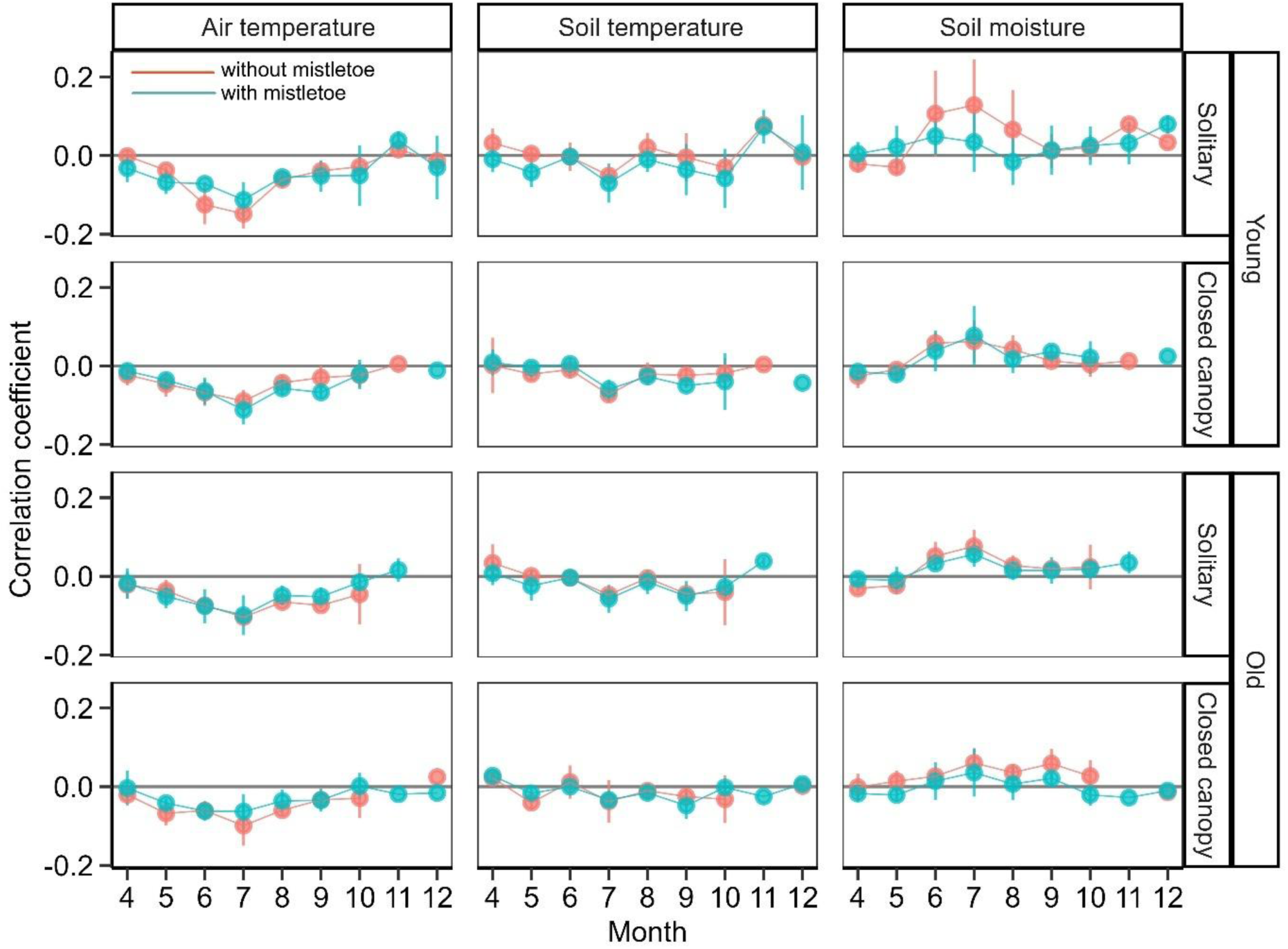
Mean (± SD) correlation coefficients between radial growth rate (15-min resolution) and (a) air temperature, (b) soil temperature, and (c) soil moisture for young and old oaks in solitary and canopy positions calculated for each calendar month with growth occurrence within 2020-2023. Blue points and line indicate trees with mistletoe infection, red points and line show non-infected trees. Correlations were calculated for individual trees and then averaged within each category. Positive values indicate faster growth under higher temperature or moisture conditions.

### Climate Optima and Infection Effects

Relationships between growth rate and climatic optima, analysed across discrete intervals of air temperature, soil temperature, and soil moisture, revealed clear non-linear responses (Fig. 5). Growth rates peaked at moderate air temperatures (10–18 °C) and corresponding moderate soil temperatures, confirming these ranges as optimal for oak radial growth. Responses of growth to soil moisture availability were rather flat.

**Figure 5.**
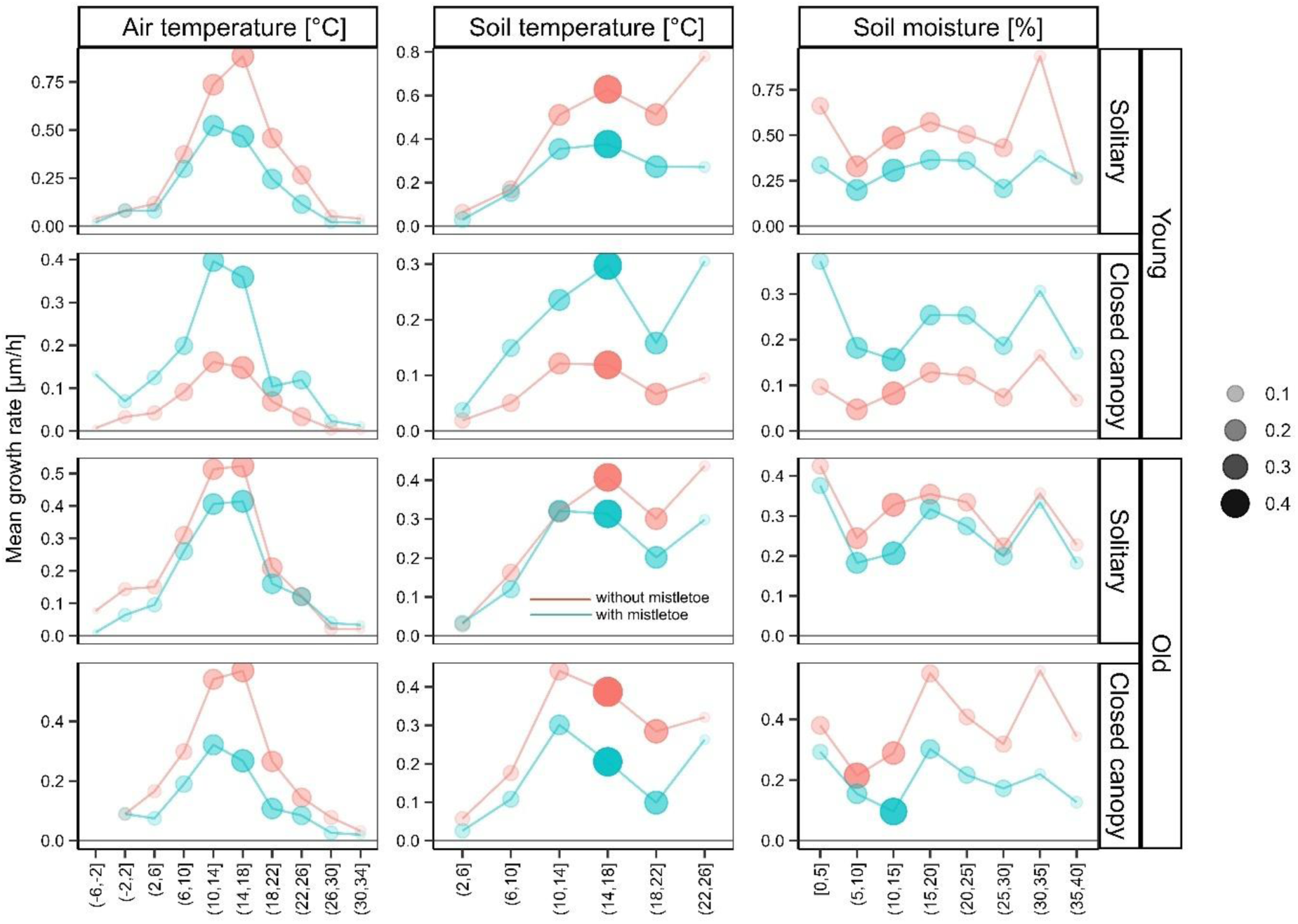
Mean radial growth rate of oak trees in relation to air and soil temperature and soil moisture across infection status (red = without mistletoe, blue = with mistletoe), age class, and canopy position. Data are binned into discrete climatic intervals; point size and its transparency reflect the relative frequency of each condition during the study period. Only bins with a relative frequency higher than 0.01 are shown. Note the different scales of the y-axes. Growth rates peak at moderate air (10–18 °C) and soil temperatures, indicating optimal climatic conditions for oak growth, while the response of growth to soil moisture is non-systematic with frequent intervals. Differences between infected and non-infected trees are generally most pronounced near climatic optima, whereas under sub-optimal conditions growth rates converge.

Differences between infected and non-infected trees were generally most evident near climatic optima, when absolute growth rates were highest. For example, in young solitary trees, non-infected individuals often outperformed infected ones at optimal temperatures and across full range of soil moisture. In contrast, under suboptimal conditions (low or high temperatures) the two groups grew at similar rates. A notable exception occurred in young canopy trees, where infected individuals matched or exceeded non-infected growth near optima of both temperatures and across a full range of soil moistures.

These results indicate that mistletoe effects on host growth are not uniform but instead are most pronounced under favourable climatic conditions, consistent with the idea that resource competition between host and parasite intensifies when both have high physiological activity. However, category-specific deviations from this pattern, such as the reversed trends in young canopy trees, highlight the complexity of host–parasite interactions in heterogeneous microhabitats.

### Joint Effects of Temperature and Soil Moisture

The combined analysis of air temperature and soil moisture revealed that oak radial growth was highest under moderate temperatures (10–18 °C) and relatively high soil moisture levels (25-30 %, Fig. 6). Within this optimal range, mean growth rates frequently exceeded 1.0 µm h⁻¹, with maximum values approaching 1.5 µm h⁻¹. Growth declined sharply at both low temperatures (< 6 °C) and high temperatures (> 22 °C), regardless of soil moisture availability. The frequency distribution of climatic conditions, represented by point size, indicated that optimal combinations of temperature and moisture were relatively common during the measurement period, especially in late spring and early summer. However, the most common conditions for our site, characterized by air temperature 14-22 °C and soil moisture 5-15 %, were rather suboptimal for tree growth, resulting in relatively low growth rates (< 0.5 µm h⁻¹). Patterns were statistically similar between infected and non-infected trees (Mantel R = 0.93, p<0.01), with both groups showing the same climatic optima. However, in optimal conditions (14–18 °C, 25-35 %), non-infected trees exhibited slightly higher mean growth rates than infected ones. This difference diminished under suboptimal climatic combinations, suggesting that the competitive effects of mistletoe on host growth are most apparent when environmental conditions support maximum physiological activity.

**Figure 6.**
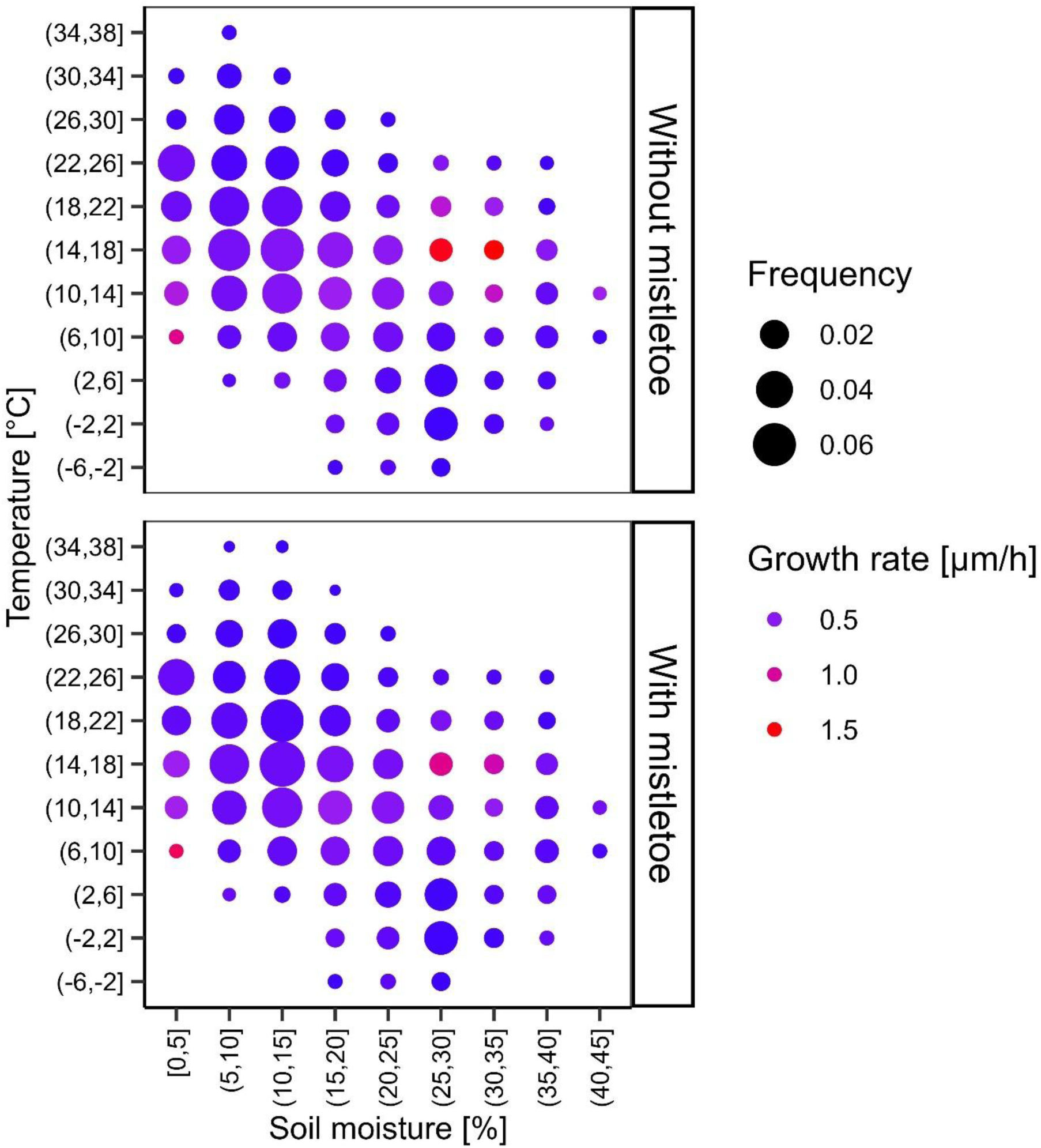
Mean radial growth rate (µm h⁻¹) of oak trees for specific combinations of air temperature (y-axis) and soil moisture (x-axis), shown separately for trees without mistletoe (top) and with mistletoe (bottom). Point size indicates the frequency of occurrence of each climatic combination during the study period, and point colour represents mean growth rate (purple = low, red = high). Only points with relative frequency higher than 0.001 are shown.

## Discussion

Our multi-year, high-resolution monitoring of oak radial growth provides new evidence that the effects of European mistletoe (*Loranthus europaeus*) on host performance are highly context-dependent, varying with tree age, canopy position, climatic conditions, and their interaction. While mistletoe infection rarely altered the phenology of radial growth, it frequently influenced growth magnitude, with the largest differences between infected and non-infected trees occurring under conditions near the climatic optimum for oak growth. These findings align with host–parasite theory for hemiparasitic plants (Těšitel, 2016), which predicts that competition for resources is most intense when both host and parasite exhibit high physiological activity, and that under stressful conditions the parasite may rely more on its own photosynthetic capacity. They are also consistent with evidence from ring-porous oaks that cambial activity and earlywood formation begin before or around leaf flush, when transpiration is still low, implying that infection contrasts may strengthen later in the season as leaf area and transpirational demand increase (Pérez-de-Lis et al., 2016; Lavrič et al., 2017; Szatniewska et al., 2022; Puchałka et al., 2017).

### Context dependence and synthesis with previous work

Infection effects in our study were strongest in young, solitary trees under favourable weather, indicating that high host vigour and full light exposure amplify resource competition. This supports a state-dependent view of hemiparasite impacts: where vigour and light are high, infection chiefly reduces growth magnitude rather than shifting phenology. That pattern is consistent with systems in which the balance of competition and facilitation varies with host size, stand density and stress (Queijeiro-Bolaños et al., 2016). Under open, high-light conditions, elevated vapour-pressure deficit together with the parasite’s sustained transpiration demand intensify hydraulic competition, so penalties become most apparent when climate approaches the host’s operational optimum. At the opposite end of the gradient, recurrent drought can push interactions past an anatomical–hydraulic threshold; in *Quercus brantii*, post-2011 droughts coincide with smaller vessels, higher vessel density and persistently reduced growth in infected trees (Oladi et al., 2025). Taken together, these lines of evidence and our results indicate acute penalties near climatic optima and chronically entrenched penalties once drought reconfigures xylem structure.

Physiological contrasts between infected and non-infected oaks observed elsewhere map closely onto the growth patterns we detected. During a drought year, Kubov et al. (2020) found that infested Q. petraea reduced stomatal conductance and transpiration (to ∼66% of non-infested trees), consistent with a more conservative, isohydric-like response of the host, while mistletoe maintained much higher stomatal conductance and transpiration; the mistletoe: host transpiration ratio was ∼3.3. This asymmetry was matched on the carbon side: non-infested oaks assimilated nearly twice as much CO₂ as either infested oaks or the mistletoe, and intrinsic water-use efficiency (WUEᵢ) was highest in non-infested trees and lowest in mistletoe; infested trees also showed higher intercellular CO₂ despite lower conductance, indicating diffusion limitation. These physiological shifts are congruent with our finding that infection suppresses growth magnitude most where climatic conditions would otherwise permit rapid growth, and they offer a mechanistic explanation for the weaker mid-summer moisture responsiveness of infected young solitaries we observed.

Stand structure and host traits further shape this pattern. Infection is most likely on large, less-competed trees and spreads over short neighbourhood distances (≈ 5 m), concentrating *Loranthus europaeus* on solitary, well-lit oaks (Matula et al., 2015), a distribution consistent with bird-mediated dispersal and the use of free-standing trees as perches by mistletoe-dispersing thrushes. In such crowns, high resource flux allows parasite demand to translate into measurable growth loss, matching the stronger infection effects we observed in solitaires. By contrast, shaded young canopy trees sometimes matched or exceeded the growth of uninfected counterparts, an outcome compatible with trait-mediated, non-trophic effects whereby mistletoes modify host architecture or physiology and thereby alter local light and water relations (Lázaro-González et al., 2019). Context dependence of this kind aligns with broader evidence in oaks that the combined influence of infestation, competition and climate depends on tree size and site conditions (Doležal et al., 2016). They also correspond with longer-term demographic and physiological studies indicating that mistletoe infection may accelerate decline in already stressed hosts (Doležal et al., 2010), while having weaker or more variable effects in vigorous individuals (Gosling et al., 2024).

A further determinant is within-parasite competition. Intraspecific competition among parasite ramets can alter realised virulence (Nabity et al., 2021). Applied to *L. europaeus*, spatial variation in clump number and size within crowns generates density-dependent draw on xylem flow, explaining why penalties in our dataset were strongest in bright, vigorous crown sectors, where many ramets remain physiologically active, and weaker under shade or stress. This perspective also clarifies the distinction, evident in Matula et al. (2015), between infection probability (driven by host size and neighbourhood structure) and infection intensity (reflecting local parasite load), and helps account for the category-specific outcomes we report across years. In combination, these mechanisms align our observation of larger growth reductions in young solitaires during favourable weather with a general framework in which host condition, stand context and parasite density jointly govern the magnitude and persistence of hemiparasite impacts.

### Climatic modulation of infection effects

Climatic conditions strongly modulated mistletoe impacts. Growth–climate relationships confirmed moderate air (10–18 °C) and soil temperatures as the optimal range for oak radial growth. Infection-related growth reductions were generally greatest in this range, particularly for solitary trees, and diminished under suboptimal conditions such as high temperatures or drought. A mechanistic link to leaf-level physiology is evident from Kubov et al. (2020): under drought, infested oaks restricted stomatal conductance and transpiration while mistletoe maintained high conductance, elevating host water limitation and depressing assimilation relative to non-infested trees. This is consistent with Bell et al. (2019) and Tamudo et al. (2021), who showed that warm or dry conditions can intensify hemiparasite effects by increasing host water stress, though in some systems the greatest negative impacts occur during favourable years when host and parasite compete at peak activity. He et al. (2021) similarly reported that host–parasite contrasts were most evident in the wet season, whereas during the driest period, infested and uninfested oaks showed similar midday water potentials. In their system, parasites maintained lower (more negative) water potentials and higher conductance only when moisture supply permitted, mirroring our largest radial-growth penalties under moderate temperatures with adequate soil water. Together, these results indicate that hemiparasite demand scales with environmental supply: when transport capacity is high, competition for water and nutrients translates directly into reduced host growth; when supply is constrained, both partners are equally limited and differences collapse.

Our climatic optima for oak growth (moderate temperatures with ample moisture) map also closely onto a VPD framework in which *Q. robur* achieves maximal growth around 12– 16 °C and vapour pressure deficit (VPD) ≤ 0.1 kPa (Tumajer et al., 2022). In this humid, growth-conducive niche, we observed the largest penalties of infection, consistent with the expectation that parasite–host competition is most consequential when VPD allows sustained cambial activity. As VPD increases or temperature deviates from its optimum, *Q. robur* growth declines sharply and becomes ‘event-driven,’ which helps explain why differences between infected and non-infected trees converge under suboptimal conditions. The stronger mid-summer moisture responsiveness of our young solitary, non-infected trees accord with this mechanism, where short moisture pulses can be exploited, infection dampens the conversion of water availability into growth. This is consistent with parasite-imposed hydraulic load observed under monsoon conditions (He et al., 2021).

Physiological evidence supports this mechanism. Escher et al. (2004) demonstrated pronounced seasonal shifts in amino compound composition of xylem sap in *Viscum album* and its hosts, reflecting changing nutrient demand and uptake patterns through the year. Walas et al. (2022) further showed that *V. album* maintains stomatal opening under drought, sustaining its own carbon gain but exacerbating host water loss. Our finding that growth suppression was most evident near climatic optima suggests that, when water and nutrient transport are maximal, parasite demand directly constrains host cambial activity. Conversely, during stressful conditions, the partial photosynthetic independence of hemiparasites (Tennakoon & Pate, 1996) may reduce the degree of competitive suppression. Superimposed on these climate effects, density-dependent interactions among parasite ramets (Nabity et al., 2021) could explain why infection penalties are most visible on well-lit, fast-growing hosts that can physiologically support many actively transpiring shoots.

Importantly, we did not detect a consistent seasonal pattern in infection effects attributable to phenological offsets between host and parasite. Although *Q. robur* initiates radial growth early in spring, before leaf flush and at low transpiration rates, differences between infected and non-infected trees were not systematically smaller in early than in late season, again consistent with the partial decoupling of cambial onset from leaf phenology in oaks (Pérez-de-Lis et al., 2016; Lavrič et al., 2017; Szatniewska et al., 2022; Puchałka et al., 2017), and suggesting that short-term temperature–moisture conditions rather than fixed phenophases primarily drive infection effects.

A notable pattern emerged in young solitary oaks: the mid-summer moisture dependence was stronger in non-infected trees than in those infected by mistletoe. This suggests that healthy young trees can capitalize on short moisture pulses to grow, whereas infection dampens the translation of moisture availability into growth. Mechanistically, mistletoe adds a persistent transpiration load and induces earlier stomatal closure and/or lower leaf–shoot hydraulic conductance in the host (Urban et al., 2012); thus, infected trees remain chronically water-limited and less responsive to inter-monthly variability in soil moisture. The effect is magnified in solitary trees where VPD, wind exposure, and radiation are higher than under the canopy, and where young trees likely have shallower rooting and smaller storage pools. Under a closed canopy, microclimatic buffering (lower VPD, cooler soils) reduces moisture stress for all trees, and infection differences are correspondingly small. This summer divergence in young solitary trees can be viewed as evidence that mistletoe infection converts growth from “resource-tracking” to “stress-limited” mode: once a chronic hydraulic/carbon drain is imposed, additional water does not proportionally increase growth.

### Implications for oak decline under climate change

Our findings highlight the importance of considering fine-scale climatic variability and host condition when predicting the effects of mistletoe on oak growth. Projected increases in temperature and altered precipitation regimes are likely to shift the frequency of conditions near the climatic optimum for oak growth. If optimal conditions become less frequent, the competitive suppression of host growth by mistletoe may diminish; however, under scenarios where favourable conditions coincide with high parasite prevalence, impacts on host productivity could be substantial. The physiological imbalances induced by mistletoe, affecting carbon allocation, water relations, and nutrient status (González de Andrés et al., 2024), are likely to interact with climate extremes in complex ways. This could exacerbate decline episodes, particularly in fragmented stands or on drought-prone sites where host recovery capacity is low.

## Conclusion

Our four-year, high-frequency records show that European mistletoe does not alter the timing of radial growth in *Quercus robur*; onset, peak and cessation were remarkably consistent across years and categories. However, it reduces growth magnitude in a strongly context-dependent way. Suppression was clearest in years and microhabitats that favoured rapid host growth, peaking near the climatic optimum (approximately 10 to 18 °C air temperature with adequate soil moisture) and especially in solitary, well-lit trees. When conditions were suboptimal (hot and dry or cold and wet), infected and non-infected trees converged in performance. Short-term relationships between climate and growth were stable across infection status; temperature effects were weakly negative during the main season, and soil moisture effects were positive. This pattern suggests that mistletoe does not change how growth tracks weather so much as it amplifies competition precisely when host physiological activity is highest. The stronger midsummer moisture responsiveness of young solitary non-infected trees further indicates that infection shifts hosts from a resource-tracking regime to a more stress-limited growth mode under exposed conditions.

## Author contributions

JD conceived the ideas and designed the methodology; JD, VL, and KK collected the data; JT and LP analysed the data; JD and JT led the writing of the manuscript.

## Acknowledgements

We thank Vitek Pejcha, Zuzana Chlumska and Thinles Chondol for their invaluable support with field sampling and assistance with data curation.

## Data Availability Statement

Data available from the Zenodo repository doi: XXX (Dolezal et al., 2025).

